# Representation of Distance and Direction of Nearby Boundaries in Retrosplenial Cortex

**DOI:** 10.1101/807453

**Authors:** Joeri B.G. van Wijngaarden, Susanne S. Babl, Hiroshi T. Ito

**Affiliations:** Max Planck Institute for Brain Research, 60438 Frankfurt am Main, Germany; Institute of Neurophysiology, Neuroscience Center, Goethe University, 60528 Frankfurt am Main, Germany

## Abstract

Borders and edges are salient and behaviourally relevant features for navigating the environment. The brain forms dedicated neural representations of environmental boundaries, which are assumed to serve as a reference for spatial coding. Here we expand this border coding network to include the retrosplenial cortex (RSC) in which we identified neurons that increase their firing near all boundaries of an arena. RSC border cells specifically encode walls, but not objects, and maintain their tuning in the absence of direct sensory detection. Unlike border cells in the medial entorhinal cortex (MEC), RSC border cells are sensitive to the animal’s direction to nearby walls located contralateral to the recorded hemisphere. Pharmacogenetic inactivation of MEC led to a disruption of RSC border coding, but not vice versa, indicating network directionality. Together these data shed light on how information about distance and direction of boundaries is generated in the brain for guiding navigation behaviour.

## Introduction

Rodents travel great distances in their natural habitat, establishing foraging paths on which they hunt and search for food. These paths often follow (natural) edges along the environment, providing safety and cloaking from their predators as opposed to exposure in open fields. When first introduced into novel experimental environments, rats show high levels of anxiety and timidity, resulting in defecation (Hall, 1934) and thigmotaxis (or “wall hugging”; Valle, 1970; Walsh & Cummins, 1976). Rats display reduced locomotion and seek out the safety of walls and corners, spending up to 98% of their initial time away outside of the centre area (Valle, 1970). It is only after extensive habituation, coupled with scattering of food for motivation, that rats are nudged to explore.

Once they enter the open space however, rodents are able to discriminate positions within the arena, allowing them to navigate to a desired location. This ability is manifested in the activity of neurons that fire at particular locations in space, such as place cells or grid cells, and population activity of place cells can distinguish nearby positions at several centimeter resolution in an open field arena (Brown, Frank, Tang, Quirk, & Wilson, 1998). It has been suggested that this ability is based on the estimation of distance and direction relative to landmarks in the environment, and previous studies have pointed to the importance here of environmental boundaries, such as walls or edges (Barry et al., 2006; O‟Keefe & Burgess, 1996). For example, a subpopulation of neurons in the medial entorhinal cortex (MEC) or the subiculum increase firing rates near the environmental boundaries, called border cells or boundary-vector cells (Lever, Burton, Jeewajee, O‟Keefe, & Burgess, 2009; Solstad, Boccara, Kropff, Moser, & Moser, 2008). The presence of dedicated representations of environmental borders in the hippocampus and parahippocampal regions implies a pivotal role of boundary information in generating accurate spatial representations in the brain. In accordance with this idea, border cells in MEC develop earlier than grid cells after birth, exhibiting adult-like firing fields at postnatal days 16-18, while grid cells still exhibit immature irregular firing fields (Bjerknes, Moser, & Moser, 2014). It has further been shown that position errors of firing fields of grid cells accumulate after the animal leaves a wall of an open-field arena, suggesting an error-correcting role of environmental boundaries for internal spatial representations.

While these previous studies have indicated a key role of environmental boundaries in the brain‟s spatial representation, it remains largely unclear how the boundary representation is generated and used in other brain regions for navigation. Recent work furthermore reported that the dorsomedial striatum contains cells that are active near the boundaries of the arena (Hinman, Chapman, & Hasselmo, 2019), leading to a question of functional relationships between these cells for boundary representations. This urges for detailed characterization and comparison of boundary coding between regions.

Here we report that a subpopulation of neurons in the retrosplenial cortex (RSC), a key brain region for navigation with reciprocal anatomical connections with MEC, increase their firing rate near environmental borders independent of wall identity. We discovered that firing of these RSC border cells is strongly modulated by the animal‟s head direction relative to the closest wall, providing local information about the animal‟s distance and direction to nearby boundaries. We explored under which environmental circumstances this information is generated by manipulating sensory and spatial cues in the environment. Furthermore, using decoding and pharmacogenetic inactivation techniques, we show the difference of boundary information as well as functional dependence between border cells in MEC and RSC, obtaining insights into the circuit organization of boundary representation in the brain.

## Results

### RSC cells fire in close proximity to the maze perimeter at specific distances

We performed electrophysiological recordings of neuronal activity in the retrosplenial cortex (Fig. 1a, Supplementary Fig. S1) of rats as they explored a squared open field arena and foraged for scattered chocolate pellets (Fig. 1b). All animals were sufficiently habituated to the environment and procedures, and actively explored the entire arena (Fig. 1c). The experimental setup was placed in the room with fixed landmarks to allow the animals to orient themselves relative to external features.

**Figure 1:**
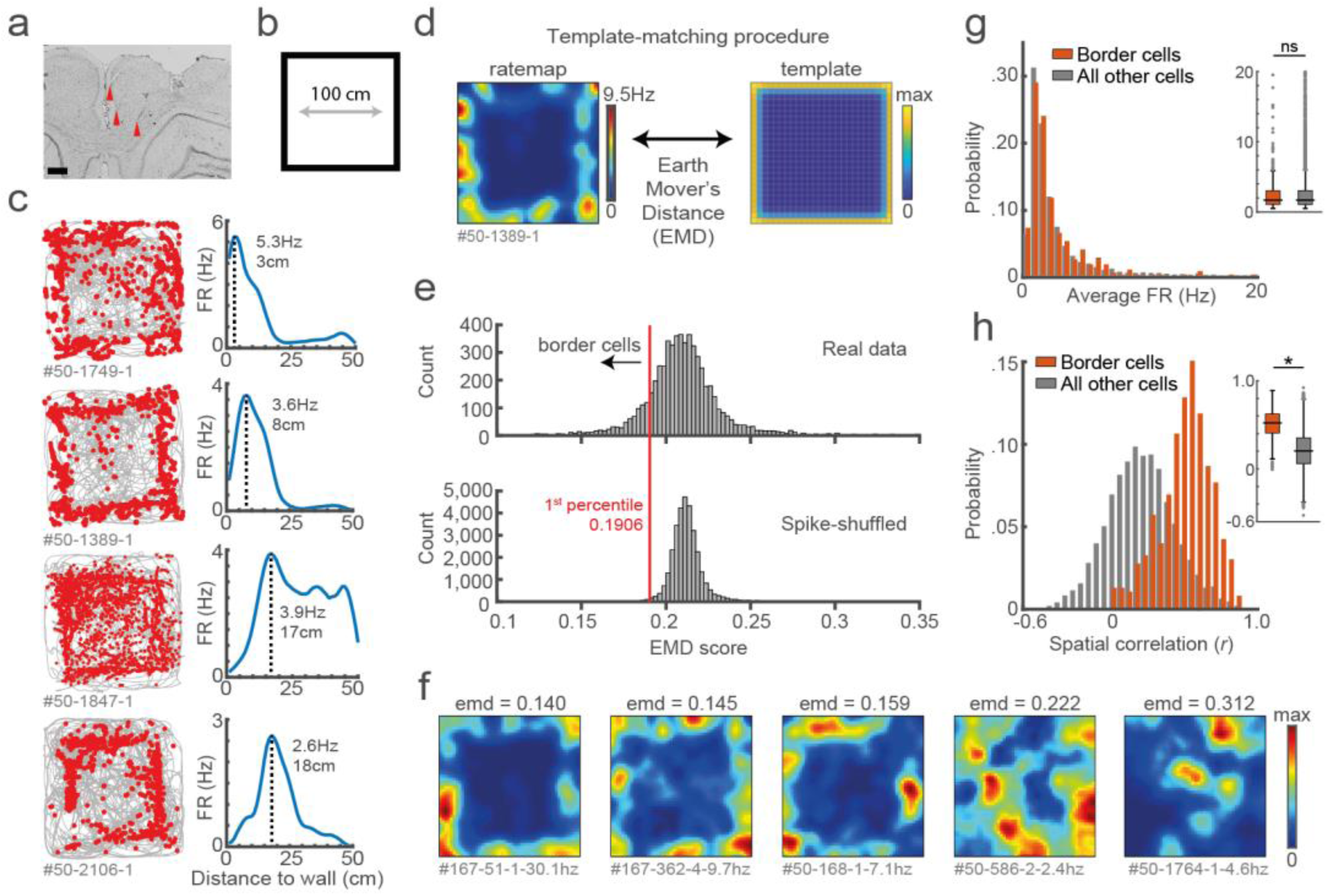
Response profiles of border cells in RSC. **(a)** Location of tetrode tracts marked with red in an example Nissl-stained coronal section. Scale bar, 500μm. **(b)** Task behaviour consisted of free exploration in a squared 1m^2^ arena. **(c)** Trajectory spike plots (left column) and distance FR plots (right column) of four example cells that fire at different distances away from the wall, relative to the closest wall at any time. Grey lines indicate the animal’s trajectory and red dots the rat’s position when a spike occurred. **(d)** A template-matching procedure was applied to classify border cells by calculating the Earth Mover’s Distance (EMD) between each cell’s spatial ratemap and an ideal template (see methods). **(e)** A cell was classified as a border cell when its EMD score was below the 1^st^ percentile of a shuffled null distribution, together with an average FR above 0.5 Hz. **(f)** Colour-coded spatial ratemaps of five example cells with different EMD scores, where warm colours indicate high firing. From left to right: three typical border cells, a non-uniform firing cell and a cell with focused firing fields. **(g)** Distribution of average FR over the entire recording day for border cells and other recorded cells. **(h)** Distribution of spatial correlations between recorded sessions for all neurons. *p < 0.05, Wilcoxon ranksum test.

We recorded the activity of 4754 RSC neurons across 8 animals (n = 75 sessions) and observed a subpopulation of cells that fired consistently at the edge of the arena (Fig. 1c). Across this subgroup there was a variety of preferred firing distances from the wall, ranging from the very near proximity up to a body-length (15-18 cm) away. Unlike traditional border cells found in MEC and Subiculum (Solstad et al., 2008; Stewart, Jeewajee, Wills, Burgess, & Lever, 2014), these border responses occurred throughout the environment on each of the four available walls. RSC border cells furthermore form multiple firing fields that are not necessarily directly connected to the wall. Typical border cell classification using the original border score (Solstad et al., 2008) identified only a small fraction of border cells in RSC, as this score is based on the occupancy of a single firing field along a wall and is strongly biased to connected bins (Supplementary Fig. S2). We thus developed a new model-based approach using a template-matching procedure to classify these border cells in RSC (Fig. 1d-1f), based on (Grossberger, Battaglia, & Vinck, 2018).

This method uses two-dimensional (2D) information of the firing rate maps and builds on the assumption that border cells have their spikes concentrated at the entire outer ring of the arena, incorporating geometric information into the classification procedure. The dissimilarity between a cell‟s spatial firing rate map and a “border” template (Fig. 1d, 1e) was assessed by the algorithm based on the Earth Mover’s Distance (Hitchcock, 1941; Rubner, Tomasi, & Guibas, 1998) (EMD; see methods), a distance metric from the mathematical theory of optimal transport. While the metric is sensitive to a change in the geometric shape of rate maps, it is robust to small variations of preferred firing distances or pixel-by-pixel jittering, giving a single tuning metric that can assess changes in the cell’s firing as a function of experimental manipulations.

Border cells were defined as stable cells with a low dissimilarity EMD score below the 1^st^ percentile of a spike-shuffled null distribution of 0.191, and an average firing rate above 0.5 Hz. In total 407 out of 4754 RSC cells (8.6%) passed this criterion (Fig. 1e, 1f). Selected border cells had a similar distribution of average firing rates compared to other recorded cells (border cells: FR = 1.70 ± 0.20 Hz, others: FR = 1.68 ± 0.08 Hz; Wilcoxon ranksum test, z = 0.024, p = 0.981; Fig. 1g), but had significantly higher spatial correlations between the first and last recording sessions (border cells: *r* = 0.52 ± 0.01, others: *r* = 0.20 ± 0.003; Wilcoxon ranksum test: z = −23.46, p = 1.15 × 10^−121^; Fig. 1h).

### Border cells form new firing fields nearby added walls but not to objects

We next asked if the firing of these border cells is limited to walls, or whether these cells also encode information about other features of the environment (e.g. local cues or objects (Hoydal, Skytoen, Andersson, Moser, & Moser, 2019; Jacob et al., 2017)). Our first manipulation was to temporarily add an additional wall, protruding from one side into the centre of the maze (Fig. 2a, 2b). Border cells formed new firing fields around the added walls accordingly, as their firing rate inside a region-of-interest (ROI) around the wall increased significantly in the added wall sessions (Regular: FR = 1.19 ± 0.13 Hz; Added wall: FR = 1.58 ± 0.21 Hz; Wilcoxon signed rank test: z = −2.67, p = 0.0076; n = 42 border cells; Fig. 2c). This was accompanied by a sharp drop in spatial correlations between ratemaps of regular versus added wall sessions (Reg-Reg: *r* = 0.51 ± 0.004, Reg-Wall: r = 0.25 ± 0.006; Wilcoxon signed rank test: z = 4.43, p = 9.31 × 10^−6^; Bonferroni-corrected α = 0.025; Fig. 2d), while correlations remained high when comparing within session types (Wall-Wall: r = 0.47 ± 0.005; Wilcoxon signed rank test with Reg-Reg correlation: z = 0.63, p = 0.53; Bonferroni-corrected α = 0.025; Fig. 2d). The EMD metric furthermore showed a significant increase in dissimilarity between ratemaps of these added wall sessions and the original border template (EMD score template 1: R1, 0.176 ± 0.002, W1, 0.207 ± 0.005, W2, 0.215 ± 0.005, R2, 0.179 ± 0.002; Friedman test: *X*^2^(3) = 77.9, p = 8.6 × 10^−17^; Post-hoc Wilcoxon signed rank test: R1-W1, z = −5.35, p = 9.0 × 10^−8^, R1-W2, z = −5.58, p = 2.37 × 10^−8^, R1-R2, z = −1.27, p = 0.20; Bonferroni-corrected α = 0.017; Fig. 2e). In contrast, the dissimilarity between the same ratemaps and an “added wall” template decreased significantly (EMD score template 2: R1, 0.145 ± 0.002, W1, 0.132 ± 0.003, W2, 0.135 ± 0.004, R2, 0.153 ± 0.003; Friedman test: *X*^2^(3) = 33.7, p = 2.3 × 10^−7^; Post-hoc Wilcoxon signed rank test: R1-W1, z = −3.89, p = 9.8 × 10^−5^, R1-W2, z = 2.59, p = 0.0095, R1-R2, z = − 2.22, p = 0.027; Bonferroni-corrected α = 0.017; Fig. 2e), confirming that border cells indeed encode wall information.

**Figure 2:**
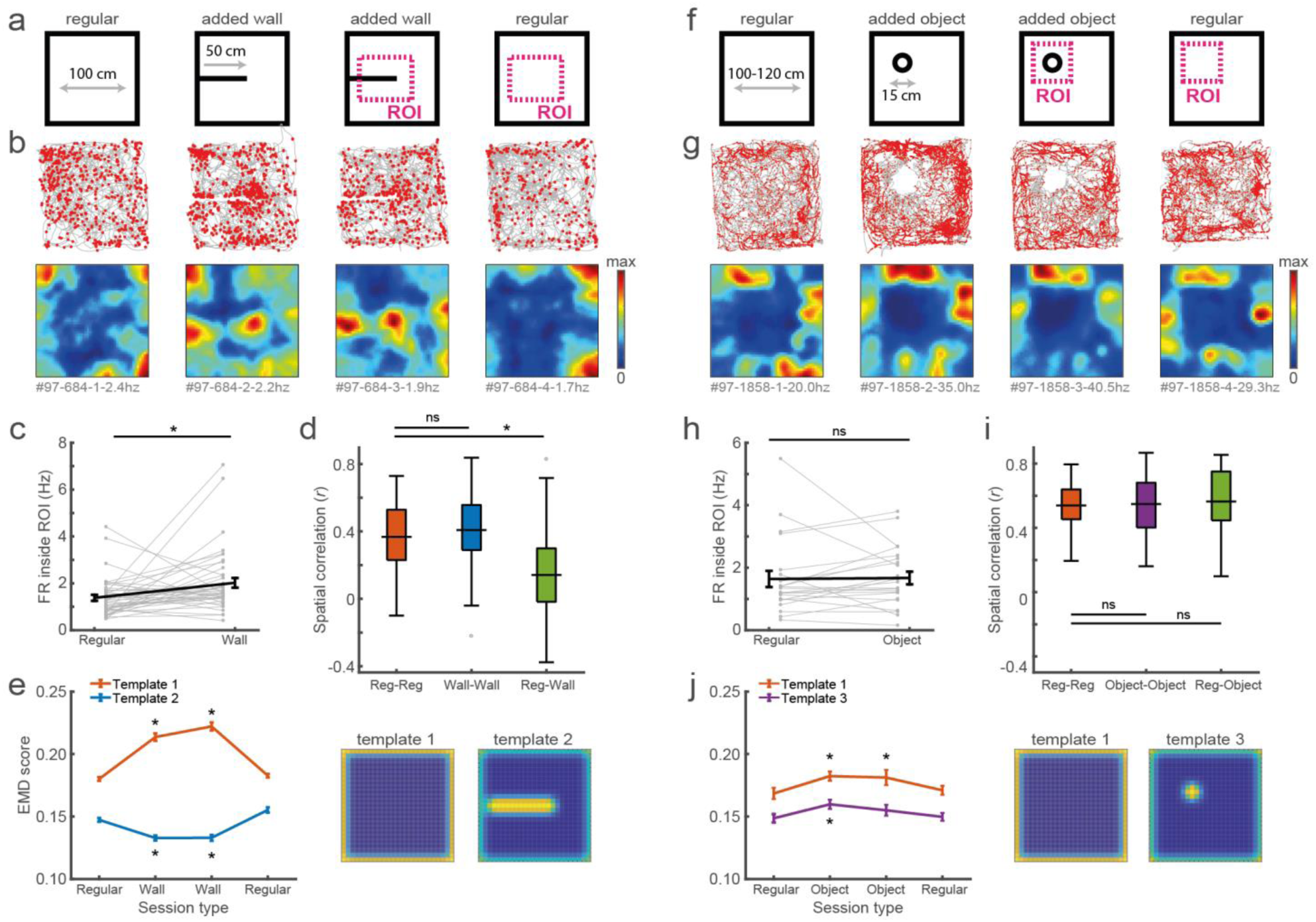
Border cells respond to new walls but not to the addition of new objects. **(a)** An additional temporary wall was added to the centre of the maze in the middle sessions. **(b)** Trajectory spike plots and spatial ratemaps of an example border cell across regular and added wall sessions during one recording day. **(c)** Border cells form new firing fields nearby the added wall, as cells significantly increase their firing rate in the region-of-interest (ROI) area around the central wall. **(d)** Spatial correlations between ratemaps of regular and wall sessions are decreased, but remain high within session type. **(e)** The original border template (#1) significantly increases in dissimilarity as cells form fields around the added wall, opposite to an added wall template (#2) which decreases in dissimilarity. **(f)** A high, non-climbable object was introduced in the north-west corner of the maze. **(g)** Trajectory spike plots and spatial ratemaps of an example border cell across regular and object sessions. **(h)** Border cells ignore the addition of objects as their FR in a ROI around the object remains unchanged between session types. **(i)** There are no significant changes in spatial correlations between the different session types. **(j)** There is a small increase in EMD scores for template 1, but objects do not elicit a response from border cells as the object template (#3) shows a similar increase. *p < 0.05 (Bonferroni correction for multiple comparisons), Wilcoxon signed rank test.

To investigate generalization to other environmental features we further added additional objects to the arena and tested the specificity of border responses to the spatial layout (Fig. 2f, 2g). Contrary to an added wall, RSC border cells maintained tuning only to the outer walls and did not fire whenever objects were inside their receptive field (Regular: FR = 1.39 ± 0.26 Hz; Added object: FR = 1.44 ± 0.20 Hz; Wilcoxon signed rank test: z = −0.57, p = 0.57; n = 23 border cells; Fig. 2h). There were no significant changes when comparing spatial correlations across session types (Reg-Reg: r = 0.54 ± 0.007, Reg-Object: r = 0.63 ± 0.009; Wilcoxon signed rank test: z = −1.41, p = 0.16; Object-Object: r = 0.55 ± 0.008; Wilcoxon signed rank test with Reg-Reg correlation: z = −0.51, p = 0.61; Bonferroni-corrected α = 0.025; Fig. 2i). EMD analyses showed a minor but significant increase in dissimilarity to the border template in the object sessions (EMD score template 1: R1, 0.169 ± 0.005, O1, 0.182 ± 0.004, O2, 0.181 ± 0.006, R2, 0.171 ± 0.004; Friedman test: *X*^2^(3) = 14.7, p = 0.002; Post-hoc Wilcoxon signed rank test: R1-O1, z = −2.71, p = 0.007, R1-O2, z = −2.80, p = 0.005, R1-R2, z = −0.79, p = 0.43; Bonferroni-corrected α = 0.017; Fig. 2j), indicating small changes in the ratemaps of the object sessions. The cells did not form new firing fields around the object however, as fitting an “object” template led to a similar increase rather than decrease in dissimilarity (EMD score template 3: R1, 0.149 ± 0.003, O1, 0.160 ± 0.004, O2, 0.155 ± 0.004, R2, 0.150 ± 0.003; Friedman test: *X*^2^(3) = 12.4, p = 0.006; Post-hoc Wilcoxon signed rank test: R1-O1, z = −2.65, p = 0.008, R1-O2, z = −1.55, p = 0.12, R1-R2, z = −0.30, p = 0.76; Bonferroni-corrected α = 0.017; Fig. 2j). Taken together these results imply that RSC border cells encode information that is specific to boundaries of the spatial layout where cell responses differentiate between the types of added features.

### Border cells retain their tuning in darkness or to an edge without a wall

One way for border cells to compute information of boundaries is through direct sensory detection of the walls, for example by whisking or visual observation (Raudies & Hasselmo, 2012). We next investigated the importance of direct sensory input on border tuning by removing either visual or somatosensory information of the boundary (Fig. 3a, 3e). We first recorded in complete darkness using an infrared position tracking system, but observed no significant changes in EMD dissimilarity scores across the sessions (EMD score template 1: R1, 0.183 ± 0.001, D1, 0.185 ± 0.003, D2, 0.177 ± 0.003, R2, 0.182 ± 0.002; Friedman test, *X*^2^(3) = 1.23, p = 0.75; n = 21 border cells; Fig. 3b, 3d). There were also no changes across spatial correlations between different session types (Reg-Reg: r = 0.42 ± 0.007, Reg-Dark: r = 0.38 ± 0.007; Wilcoxon signed rank test z = 0.61, p = 0.54; Dark-Dark: r = 0.42 ± 0.01, Wilcoxon signed rank test with Reg-Reg correlation, z = 1.20, p = 0.23; Bonferroni-corrected α = 0.025; Fig. 3c), indicating that activity is not generated solely through visual sensory input.

**Figure 3:**
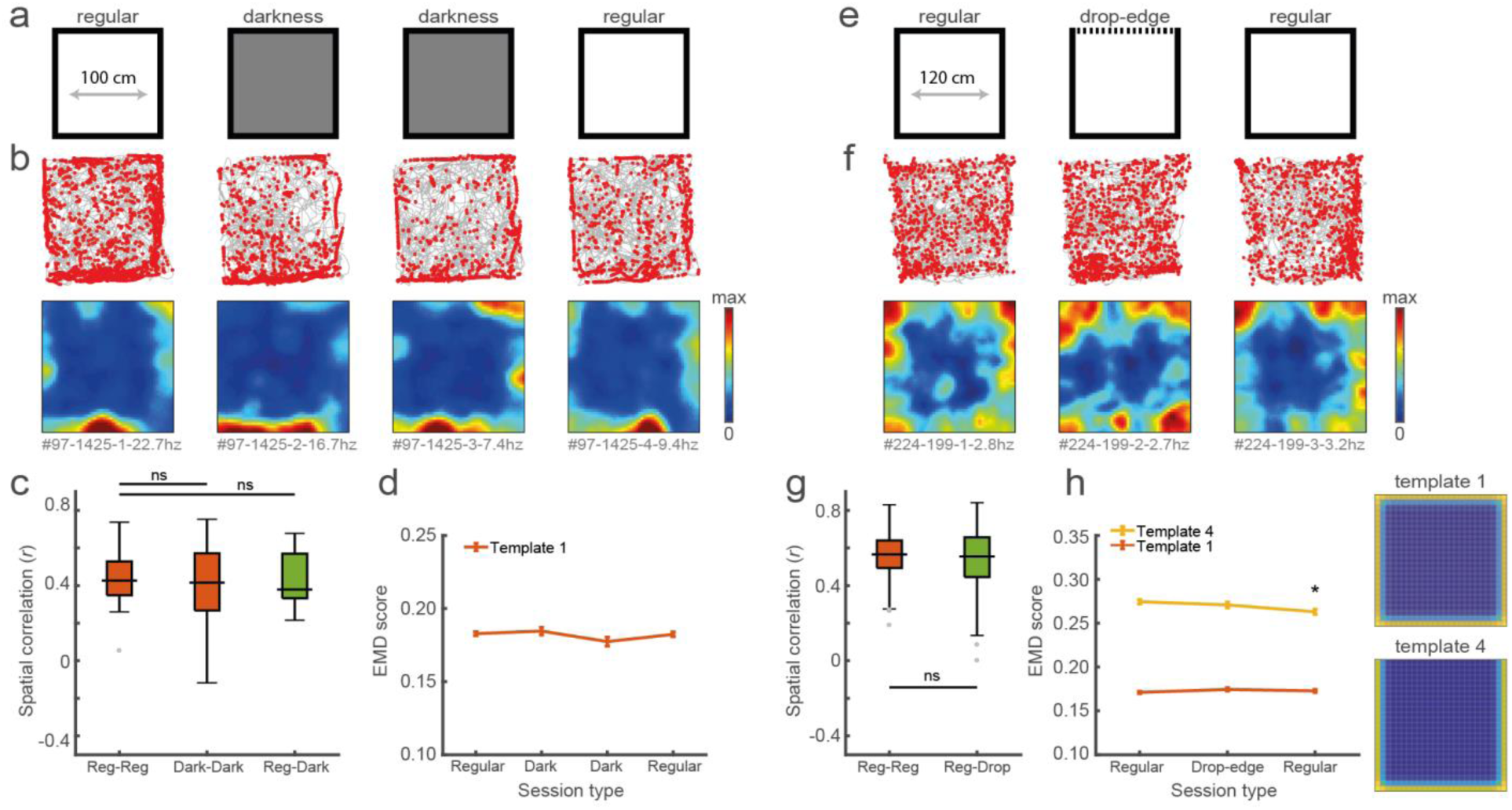
Removing direct sensory detection of walls does not alter the border cell’s activity near boundaries. **(a)** Recordings were performed in complete darkness for the middle sessions, and animals were tracked in the non-visible infrared spectrum. **(b)** Trajectory spike plots and spatial ratemaps of an example border cell recorded in light and dark conditions. **(c)** Spatial correlations between ratemaps of regular and dark sessions remain high, indicating border cells still fire nearby boundaries in darkness. **(d)** There were no changes in EMD scores with template 1, confirming that cells maintain their tuning to the outer walls without direct visual detection. **(e)** One of the outer walls was removed, leaving only a drop-edge on one side to confine the arena. **(f)** Trajectory spike plots and spatial ratemaps of an example border cell across recording sessions. **(g)** There are no significant changes in spatial correlations across session types. **(h)** Spatial ratemaps remain unchanged across session type, with no relevant changes in EMD scores for either template. * p <0.05 (Bonferroni correction for multiple comparisons), Wilcoxon signed rank test.

Similarly, we removed one of the outer walls that left a drop-edge above the floor, limiting movement of the animal in the absence of direct somatosensory information of a physical barrier (Fig. 3e). Again there were no major changes in EMD dissimilarity scores of the original border template for the regular versus drop-edge sessions (EMD score template 1: R1, 0.171 ± 0.001, Drop, 0.174 ± 0.002, R2, 0.173 ± 0.002; Friedman test: *X*^2^(2) = 7.0, p = 0.03; Post-hoc Wilcoxon signed rank test: R1-Drop, z = −2.04, p = 0.041, R1-R2, z = −2.03, p = 0.041; Bonferroni-corrected α = 0.025; n = 78 border cells; Fig. 3f, 3h). We also observed no relevant changes in dissimilarity for a “drop-edge” template across all sessions, besides a small though significant drop in the final regular session (EMD score template 4: R1, 0.274 ± 0.002, Drop, 0.271 ± 0.003, R2, 0.263 ± 0.003; Friedman test: *X*^2^(2) = 17.5, p = 0.0002; Post-hoc Wilcoxon signed rank test: R1-Drop, z = 0.76, p = 0.44, R1-R2, z = 4.14, p = 3.45 × 10^−5^; Bonferroni-corrected α = 0.025; Fig. 3f, 3h), indicating that RSC border cells do not change their firing properties alongside the drop-edge compared to a physical wall, in a similar manner as border cells in MEC (Solstad et al., 2008). This is supported by stable spatial correlations across session type comparisons (Reg-Reg: r = 0.57 ± 0.002, Reg-Drop: r = 0.55 ± 0.002; Wilcoxon signed rank test: z = 0.60, p = 0.55; Fig. 3g). These results suggest that neural activity of RSC border cells is not driven by pure sensory detection of boundaries, as cells are unaffected by the removal of unimodal sensory input.

### RSC cells have a biased directional tuning to boundaries in the contralateral side of the recorded hemisphere

Recent reports pointed to egocentric anchoring of spatial representations to environmental features such as the maze centre or walls (Hinman et al., 2019; LaChance, Todd, & Taube, 2019). RSC border cells described here have a similar direction tuning, where spikes that occur in close proximity to a wall are constraint by specific directions of the animal relative to the boundary (Fig. 4a). Projecting this trajectory data onto new body-centric axes, where coordinates indicate distance and direction of the nearest wall relative to the animal, indeed shows that cells fire predominantly whenever the wall occupies proximal space on the contra-lateral side of the recorded hemisphere (Fig. 4b, 4c).

**Figure 4:**
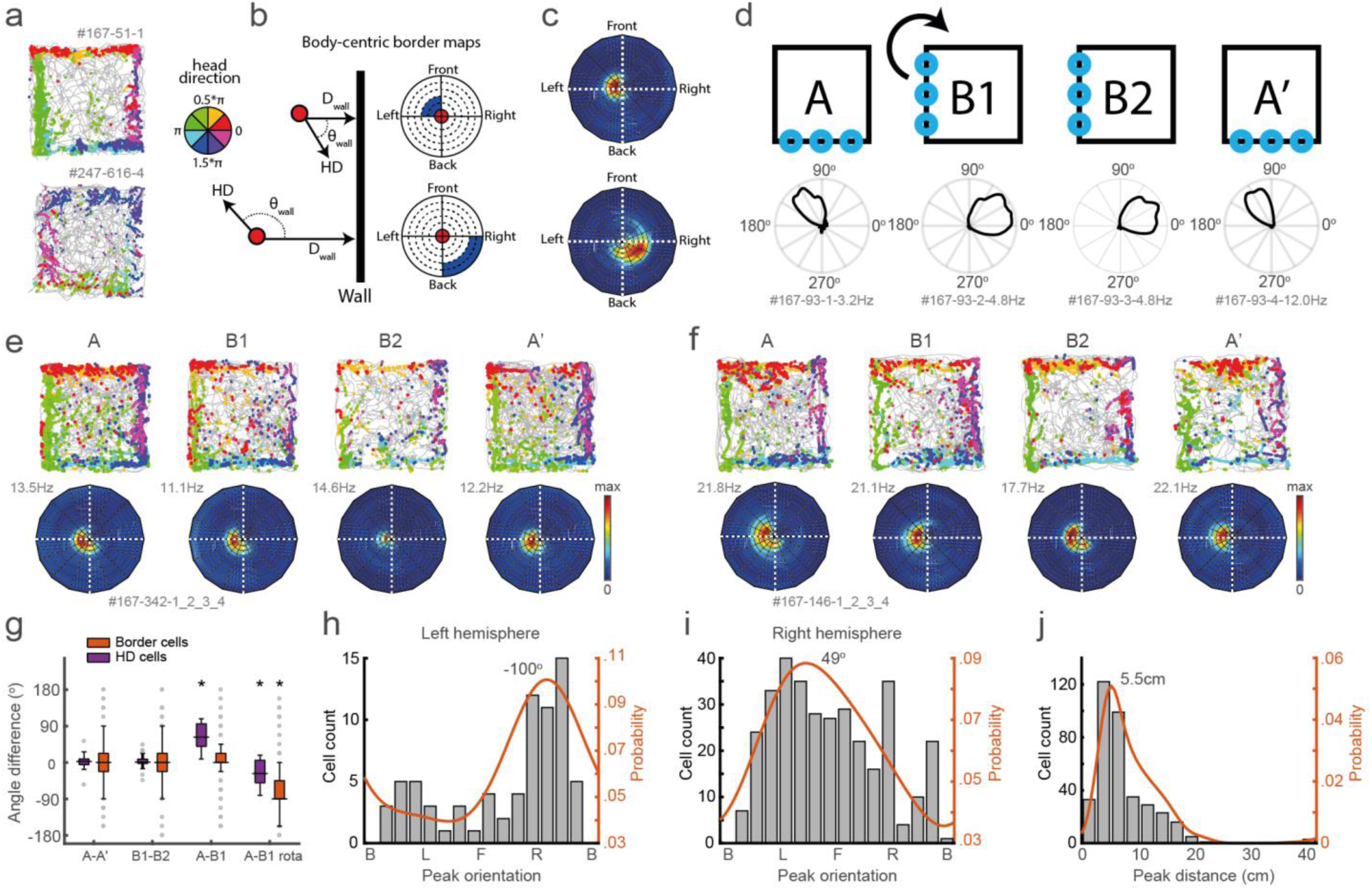
Border responses have narrow directional constraints and are biased to the contra-lateral hemisphere. **(a)** Example trajectory spike plots with spike locations colour-coded according to the direction of the animal. Most spikes alongside a wall occur only when the animal is in a narrow range of directions. Top: recorded in right hemisphere. Bottom: recorded in left hemisphere. **(b)** Trajectory data is projected onto new body-centric border maps, where coordinates indicate the distance (D_wall_) and direction (θ_wall_) of the closest wall relative to the animal’s position and head direction (HD) respectively. **(c)** Ratemaps in this border space for the same example cells shown in (a). **(d)** Top: prominent blue landmark LEDs were placed on one wall, and the entire experimental set-up was rotated 90° clockwise in the middle sessions. Bottom: example HD cell showing its tuning shifted accordingly. **(e-f)** Two example cells with trajectory spike plots and border ratemaps showing egocentric border tuning is stable across rotation sessions. **(g)** Comparison of shifts in direction tuning for head direction and border cells across the different sessions. **(h)** Preferred directional tuning of all border cells recorded in the left hemisphere, with 0° being in front of the animal. **(i)** Same as (h), but now for all border cells recorded in the right hemisphere. **(j)** Preferred distance tuning of all border cells. * p <0.05 (Bonferroni correction for multiple comparisons), Wilcoxon signed rank test.

We sought to establish whether this egocentric constraint was imposed by the head direction signal, as RSC receives inputs from the anterior limbic system that is a major source of head direction signals, and a subpopulation of RSC cells are tuned to allocentric head direction (Chen, Lin, Green, Barnes, & McNaughton, 1994; Mitchell, Czajkowski, Zhang, Jeffery, & Nelson, 2018). If the boundary representation of RSC border cells is driven by internally generated global direction signals, realignment of the head direction cells may affect the preferred tuning direction of RSC border cells. In order to manipulate the tuning of head direction cells, four blue landmark LEDs were placed on one side of the maze while all other sensory cues were kept invariant across the environment. The entire experimental setup was then rotated 90° clockwise in the middle sessions (Fig. 4d). As a result, all allocentric head direction (HD) cells rotated their tuning curves accordingly, although not a full 90° (A-A’: median shift = 2.6°, z = 1.23, p = 0.23; B1-B2: median shift = 0.8°, z = 0.61, p = 0.54; A-B1: median shift = 62.9°, z = 4.62, p = 3.8 × 10^−6^; A-B1 rotated: median shift = −27.3°, z = − 3.07, p = 0.002; Wilcoxon signed rank test; Bonferroni-corrected α = 0.013; n = 28 HD cells; Fig. 4g). The direction tuning of border cells in contrast remained unchanged (examples in Fig. 4e, 4f; A-A‟: median shift = 0°, z = 0.085, p = 0.93; B1-B2: median shift = 0°, z = −0.85, p = 0.40; A-B1: median shift = 0°, z = 1.61, p = 0.11; A-B1 rotated: median shift = −79°, z = − 3.95, p = 7.7 × 10^−5^; Wilcoxon signed rank test; Bonferroni-corrected α = 0.013; n = 46 border cells; Fig. 4e-4g). This result indicates that the direction tuning of RSC border cells is either generated by local place and direction information independent of allocentric head direction cells, or is dependent on the integration of tightly-bound allocentric position and head-direction coding that rotated together.

Across the population, border cells were tuned predominantly to the very near proximity (main peak at 5.5 cm; Fig. 4j), although some cells had fields at extended distances up to 20 cm away from the wall. Border cells showed a similar disproportionately biased distribution of preferred directions, dependent on the hemisphere where cells were recorded (Left hemisphere: mean direction = −102.9°, z = 10.11, p = 3.0 × 10^−5^; Right hemisphere: mean direction = 32.0°, z = 32.54, p = 3.4 × 10^−15^; Rayleigh test; comparing both probability distributions: two-sample Kolmogorov-Smirnov test, p = 0.021; n = 333 border cells; Fig. 4h, 4i). The majority of border cells were tuned to the contra-lateral side of the implanted electrode (e.g. whenever the wall is on the right side while the cell is recorded in the left hemisphere), although not exclusively (Fig. 4h, 4i). This hemisphere-specific tuning bias implies that boundary representations in RSC may either be generated by direct sensory signals, or reflect the command of motor actions, in both of which this bias arises along the right-left body axis.

### Inhibition of MEC disrupts border cell activity in RSC but not vice versa

The retrosplenial cortex is known to have direct, bi-directional connections with the medial entorhinal cortex (Bethany F Jones & Witter, 2007; Ohara et al., 2018), in particular with MEC layer 5 where the majority of border cells are located (Boccara et al., 2010), although their function remains unknown. Given the presence of border cells in both RSC and MEC, albeit with different properties, it is crucial to establish the direction and extent of functional interactions between these brain regions. We thus performed electrophysiological recordings of border cells in RSC and MEC and quantified their boundary information.

Border cells in MEC are different from those in RSC by having fields attached to only one or two walls rather than all (Fig. 5a) (Solstad et al., 2008), but both populations have similar peak firing rates when the animal‟s distance and direction to the wall were in the optimal range (RSC: FR = 4.08 ± 0.50 Hz, MEC: FR = 5.39 ± 1.00 Hz, Wilcoxon ranksum test: z = 0.72, p = 0.47; Fig. 5b). We first examined whether border cells in the two regions carry similar distance information on a population level. A decoder based on support vector machines estimated the animal’s distance away from the wall using population spiking activity, and performed with high accuracy for both MEC and RSC in the lower distance range (p < 0.05 for 0-20 cm, compared with a chance level of 20%; Fig. 5c). However, decoding performance from RSC activity dropped to chance level in the higher distance range (p > 0.05 for 30-50 cm; Fig. 5c), suggesting RSC border cells mainly encode local information. This matches the firing properties of RSC border cells which have preferred distance tuning up to 20cm away from the wall (Fig. 4j). Conversely, MEC computes distance information that extends well into the arena, with decoding performance above chance-level until the maximum range of 50 cm (e.g. in the centre of the maze; p < 0.05; Fig. 5c, 5d).

**Figure 5:**
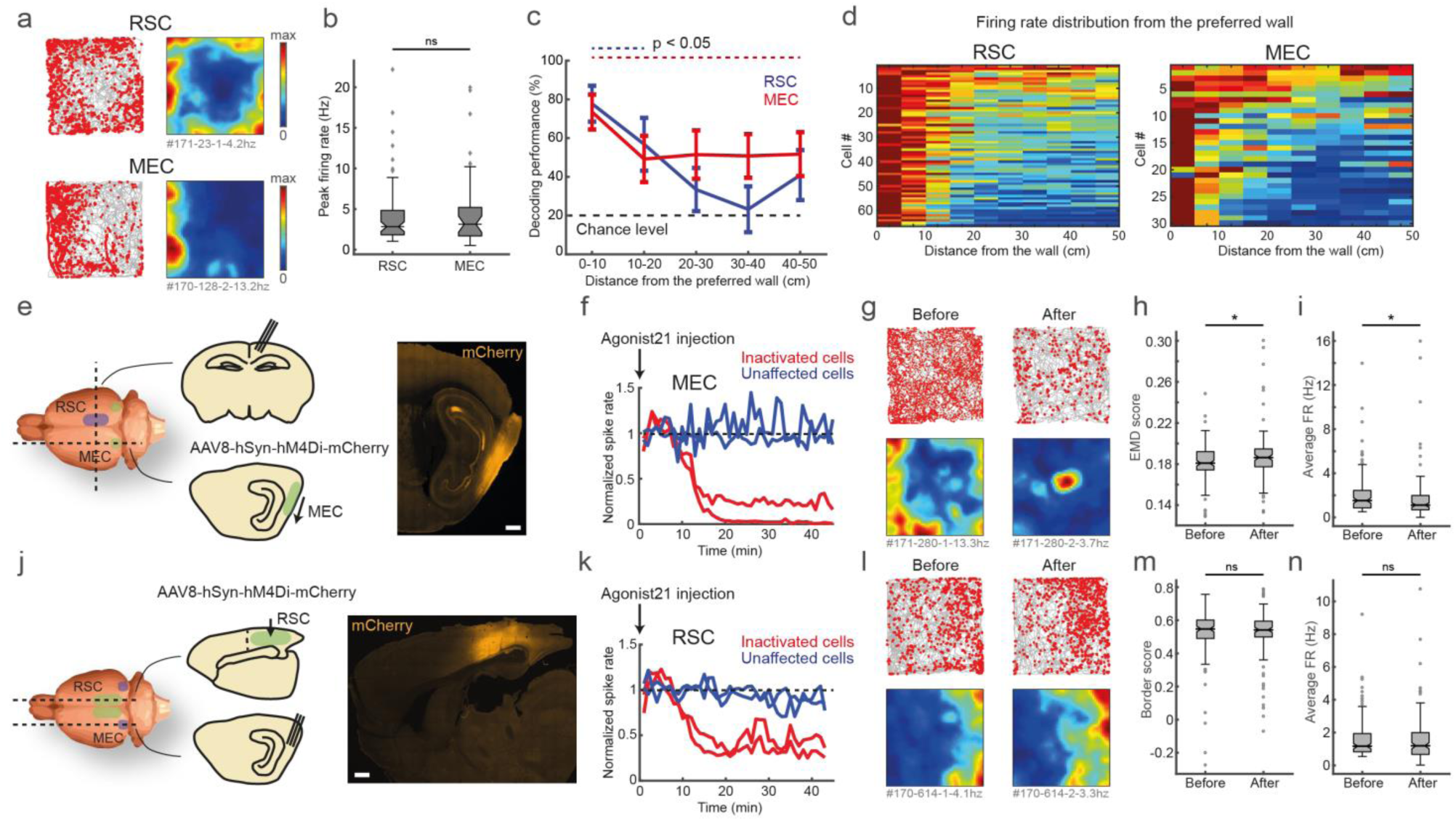
RSC border cells provide local boundary information and receive input from MEC. **(a)** Spike trajectory plots and spatial ratemaps of two border cells recorded in RSC and MEC. **(b)** Border cells in RSC and MEC have a similar distribution of peak firing rates. **(c)** A decoder using a support vector machine classifier estimated the animal’s distance to the wall based on population spiking activity. Local distance information is present in both regions, but extends further into the centre of the maze only in MEC. **(d)** Distributions of firing rate as a function of distance to the preferred wall for border cells in both brain regions. **(e)** An AAV encoding inhibitory DREADDs hM4Di was injected into MEC while tetrodes were positioned into RSC. Scale bar, 1mm. **(f)** Four tetrodes were placed locally near the virus injection site, showing affected neurons drastically decreased their activity 10-15 min after subcutaneous administration of agonist-21 (DREADDS agonist). **(g)** An example RSC border cell that is affected by MEC inhibition. **(h-i)** Border cells in RSC have decreased border tuning and lower firing rates after inhibition of MEC. **(j)** Reversed experiment, with electrophysiological recordings in MEC while the AAV was injected into RSC. Scale bar, 1mm. **(k)** Affected RSC neurons decrease their activity after administration of agonist-21. **(l)** An example MEC border cell that is unaffected by inhibition of RSC. **(m-n)** Border cells in MEC do not show any significant qualitative changes in border tuning or firing rates after RSC inhibition. * p <0.05, Wilcoxon ranksum (b) or signed rank (h-i and m-n) test.

Finally, we addressed the question of whether there is any communication between MEC and RSC in terms of encoding border information using a pharmacogenetic inactivation technique (Armbruster, Li, Pausch, Herlitze, & Roth, 2007). We first injected an AAV encoding the inhibitory DREADDs hM4Di into MEC, while simultaneously implanting a 28-tetrode hyperdrive into RSC (Fig. 5e, Supplementary Fig. 4). Subcutaneous administration of agonist-21 (DREADDs agonist) resulted in a drastic reduction of firing after 20 min for MEC cells infected with the virus (Fig. 5f). Inactivation of MEC led to a subsequent disruption of firing in a subset of RSC border cells (Fig. 5g), worsening border tuning that resulted in higher EMD scores (before: EMD score = 0.181 ± 0.002, after: EMD score = 0.186 ± 0.003; Wilcoxon signed rank test: z = −2.40, p = 0.016; n = 102 border cells; Fig. 5h) and lower overall firing rates after the manipulation (before: FR = 1.52 ± 0.20 Hz, after: FR = 1.12 ± 0.24 Hz, Wilcoxon signed rank test: z = 3.15, p = 0.0016; Fig. 5i). We next performed a reversed manipulation, injecting the virus encoding DREADDs hM4Di into RSC while recording neural activity in MEC (Fig. 5j, Supplementary Fig. S4). Administration of agonist-21 led to similar decreased activity in RSC for the infected cells (Fig. 5k), but RSC inhibition had no significant effect on MEC border cell tuning (before: border score = 0.55 ± 0.015, after: border score = 0.54 ± 0.014; Wilcoxon signed rank test: z = −0.014, p = 0.989; n = 96 border cells; Fig. 5m) or average firing rates (before: FR = 1.17 ± 0.11 Hz, after: FR = 1.19 ± 0.13 Hz; Wilcoxon signed rank test: z = 1.153, p = 0.249; Fig. 5l, 5n). Given the presence of border cells in both RSC and MEC and their bidirectional connectivity, it seems plausible that both regions are part of a broader border coding network. Our results here indeed show this to be the case, although only in one direction, suggesting that RSC border coding is partly dependent on MEC but not vice versa.

## Discussion

We have shown that a subpopulation of neurons in the RSC increase their firing rates when the animal approached the proximity of walls. We used a metric of the earth mover‟s distance to quantify the boundary coding of cells, and found that border responses are specific to boundaries that impede the movement of animals, while they are invariant to an object introduced into the maze. Border responses were maintained in complete darkness and to an environmental edge without a physical wall. These results together suggest that RSC border cells are not simply driven by local sensory cues, but likely discriminate boundaries from a global perspective of the environment.

Notably, we found that firing of RSC border cells is strongly constrained by the animal‟s head direction toward nearby boundaries, rather than to the environment, indicating body-centred or egocentric border representation. Furthermore, we assessed the spatial information provided by a population of border cells in RSC and MEC by implementing a decoding analysis and found that RSC border cells provide only local information at the wall proximity, whereas MEC border cells provide long-range distance information of a boundary. Finally, by inactivating neurons in either MEC or RSC, we found that the activity of RSC border cells is partly driven by MEC, but not vice versa. Altogether our results clarify the features of boundary representations in RSC, as well as key differences of their codes from border cells in MEC.

Anatomically, RSC locates at an interface region of the hippocampus and MEC with sensory and motor cortices (Bethany F Jones & Witter, 2007; Sugar, Witter, van Strien, & Cappaert, 2011; T van Groen & Wyss, 1990, 1992; Thomas Van Groen & Wyss, 2003). While both human patients and rodents with lesions in RSC exhibited severe impairment in navigation ability (Takahashi, Kawamura, Shiota, Kasahata, & Hirayama, 1997; Vann, Aggleton, & Maguire, 2009), the exact role of RSC has been largely unclear until recently. Several recent studies have provided clues for understanding RSC function. An fMRI study in humans demonstrated that RSC is particularly engaged in representing permanent landmarks in the environment (Auger, Mullally, & Maguire, 2012), which is consistent with the present finding of border cells as walls can serve as permanent landmarks in an open field arena, especially in the absence of local cues. On the other hand, recording studies in rats have identified several types of spatially-tuned cells in RSC, such as head-direction cells, place cells, and the cells that represent geometric features of the environment (Alexander & Nitz, 2015; Cho & Sharp, 2001; Mao, Kandler, McNaughton, & Bonin, 2017). Because of the existence of these spatially-tuned cells as well as anatomical connections, RSC has been considered an ideal brain region to implement a transformation of spatial representations between egocentric and allocentric coordinate systems (Byrne, Becker, & Burgess, 2007; Mitchell et al., 2018). The allocentric-egocentric transformation is an essential computational step for navigation because, while spatial representations in the parahippocampal regions about head direction, places, or borders, are anchored to external features of the environment (i.e. in allocentric coordinates), experiencing the world through sensory organs and executing motor plans to move through space is referenced to the actor’s body and viewpoint (i.e. in egocentric coordinates). Recent studies have reported neurons with egocentric tuning to navigational landmarks, such as the maze centre, objects, or boundaries, in brain regions including the lateral entorhinal cortex, the postrhinal cortex, and the dorsomedial striatum (Hinman et al., 2019; LaChance et al., 2019; Wang et al., 2018), and a picture is emerging of a functional network across brain regions that encode a wide-range of environmental features from a self-centred perspective.

Our findings are consistent with the RSC‟s role in coordinate transformation because both allocentric head-direction cells and egocentric border cells co-exist in RSC. The question is how such egocentric representation is generated. One possibility is that egocentric border firing is directly driven by sensory perception, such as optic flow or whisker sensation, which is egocentric in nature. However, our present results argue against this possibility as firing of RSC border cells was not affected by the absence of direct visual or somatosensory detection. Instead, our results favour the idea that RSC border cells are driven, at least in part, by MEC cells. This idea was proposed as a theoretical model (Byrne et al., 2007), in which the information about allocentric boundary locations is integrated with head-direction signals to form egocentric border representations. We found that the rotation of head-direction cells in RSC, elicited by a cue rotation of the environment, did not affect the egocentric tuning of RSC border cells, indicating that head-direction and position coding in RSC border cells must be bound and rotated together during environmental manipulations, consistent with the proposed circuit model (Byrne et al., 2007). This idea is further supported by our experiments with DREADDs-mediated activity manipulations, in which RSC border cells were significantly impaired by the inactivation of MEC, whereas RSC inactivation did not change the quality of border cording in MEC, suggesting that RSC border cells are partly dependent on MEC activity, but not likely the source of boundary information in MEC.

However, our results also clarify that RSC border cells are not necessarily a simple product of coordinate transformations from MEC cells. Our data clearly show a strong bias of tuning direction contra-lateral to the recorded hemisphere, an effect not observed in parahippocampal regions, which would indicate that a single hemisphere could transform only half of the potential behavioural space. Second, the range at which information about wall distance is present is different between MEC and RSC border cells. While RSC border cells provide local information about a nearby wall that is located less than 20 cm from the animal‟s position, border cells in MEC have extended distance information up to 50 cm (from a wall to the centre of the maze). These findings indicate that RSC border cells do not necessarily constitute an egocentric border map as a counterpart of an allocentric map in MEC.

What can be the cause of hemisphere-specific bias to boundaries in the animal‟s contralateral side, if RSC border cells are not directly driven by sensory perception? This bias may be a manifestation of the animal‟s immediate action control to the direction of an approaching wall. Collision detection and avoidance are fundamental roles of sensory-motor systems for many species of animals (Fotowat & Gabbiani, 2011), and rodents are also required to detect boundaries to avoid hitting walls or falling off edges. The boundary information in MEC and RSC may therefore be used in other brain regions to control the animal‟s next movements against walls or edges. RSC gives rise to inputs in brain regions necessary for motor control and initiation, such as premotor and motor cortices, cingulate cortex, as well as the dorsal striatum (Guo et al., 2015; B. F. Jones, Groenewegen, & Witter, 2005; Yamawaki, Radulovic, & Shepherd, 2016). A recent recording study on the dorsomedial striatum has identified a type of neurons that fire near environmental borders in a similar manner as RSC border cells do. However, their egocentric tuning is largely dependent on the animal‟s movement direction (Hinman et al., 2019), rather than head direction as in RSC border cells. RSC border cells may thus provide the downstream striatum circuits with information about the direction of approaching wall in an egocentric perspective so that animals can initiate next appropriate actions against the wall direction.

Our results thus support the idea that RSC implements a coordinate transformation of behaviourally relevant information, pointing to RSC as a key brain region linking between the brain‟s allocentric spatial representation and behaviours.

---

**Supplementary Figure S1:**
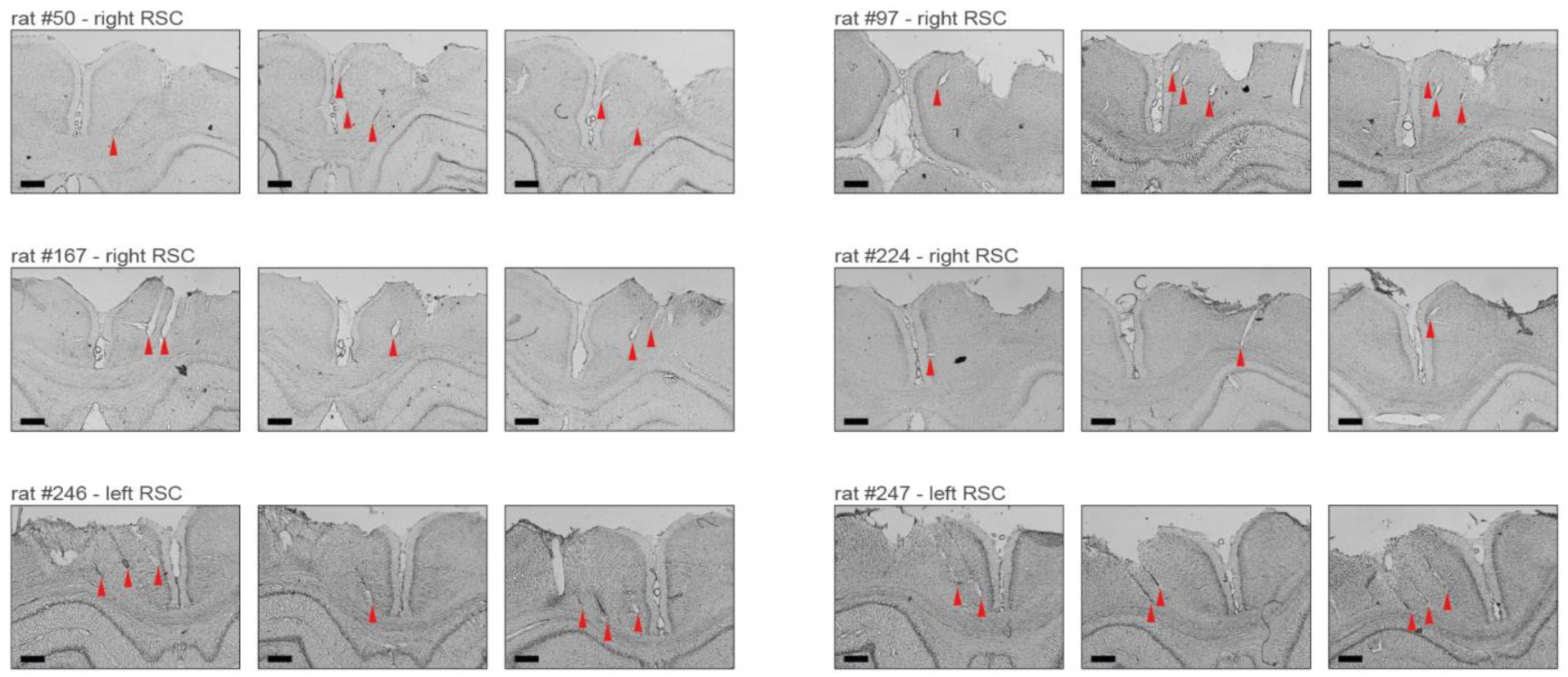
Nissl-stained coronal sections showing recording locations and tetrode tracts for all recording experiments. Shown are three typical coronal sections for each of the six animals included in the electrophysiological experiments. The top two rows include four rats (rats #50, #97, #167 and #224) with the electrode implanted in the right hemisphere, and the bottom row shows sections of two rats with a drive in the left hemisphere (rats #246 and #247). Recordings started at approximately 1mm below the surface of the cortex, and continued in a medioventral direction with a 25° angle until tetrodes reached either the midline or corpus callosum. Red triangles indicate the end of tetrode tracts. Scale bars, 500μm.

**Supplementary Figure S2:**
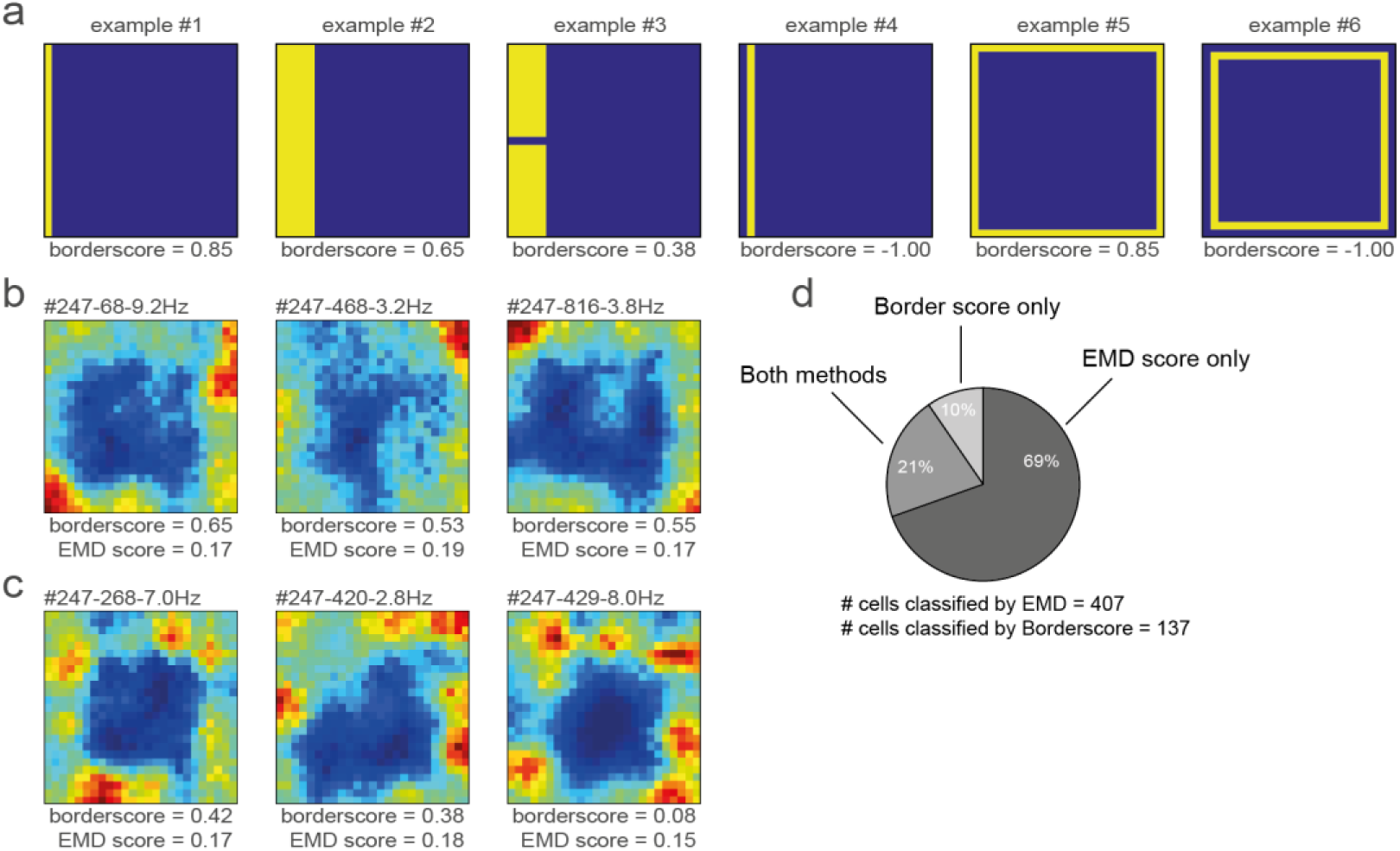
The original border score is unable to identify the majority of border cells in RSC. **(a)** Shown are 6 examples of simulated ratemaps and their associated border scores. This metric is designed to capture the coverage of a firing field alongside a single wall, and a maximal score is reached when it occupies only bins that are directly connected to the wall (#1). Extension of the field towards the centre lowers the border score (#2), as does breaking the field into two or more sub fields (#3). The algorithm is unable to calculate a border score when the firing field does not directly touch the boundary (#4). The border score does not take into account symmetry, as the maximum score on any of the four walls is selected (#5-#6). **(b)** Shown are three example RSC border cells that were classified correctly by both the border score (values above 0.5) and our EMD template matching method (values below 0.1906). **(c)** By contrast are three similar RSC border cells that were identified only by the EMD method, as these cells had low, non-significant border scores. RSC border cells tend to form firing fields that are not necessarily connected to the wall, and are often not continuous due to additional directional tuning, hence leading to low border scores. **(d)** Distribution and overlap of border cell classification using the border score and EMD methods.

**Supplementary Figure S3:**
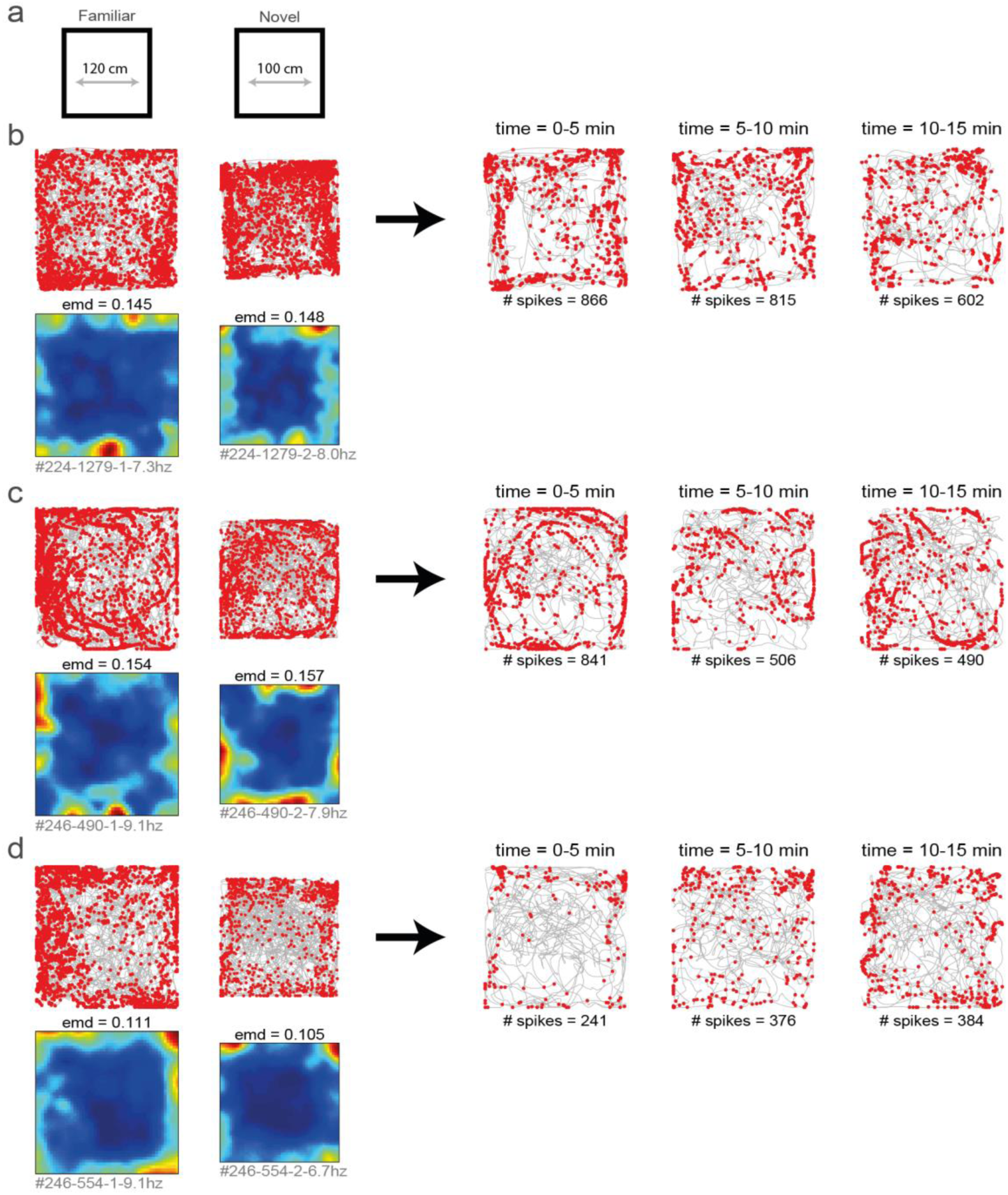
RSC border cells fire from the start in a completely novel environment. **(a)** Several experimental sessions were performed under novel conditions; animals had never visited neither this maze nor the recording room before. **(b-d)** Shown are trajectory spike plots and spatial ratemaps of three example border cells in a familiar and novel room. Shown on the left is data of the entire recorded session. On the right a subdivision of only the novel session into blocks of 5 minutes each.

**Supplementary Figure S4:**
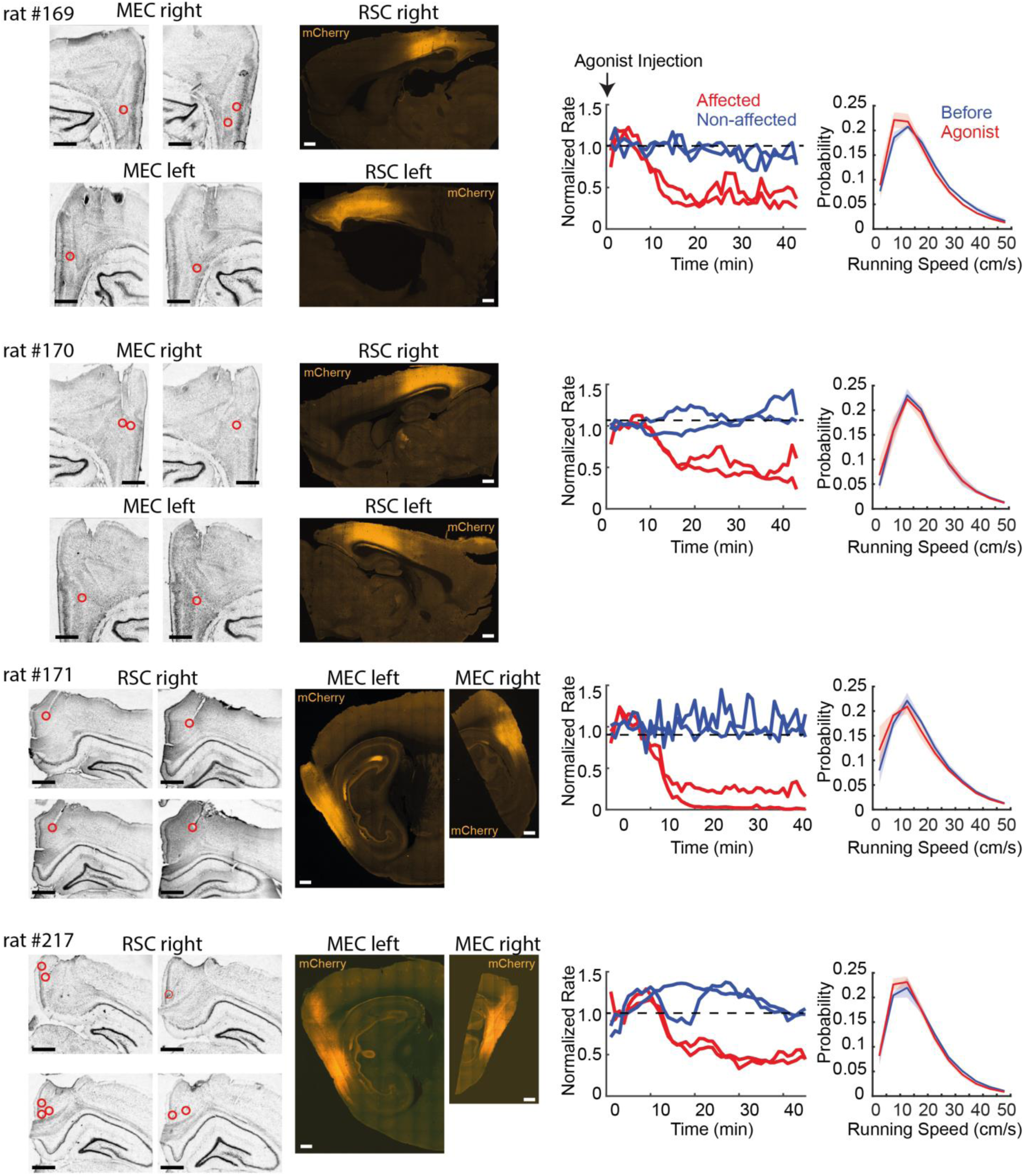
Tetrode locations and hM4Di expressions in the experiments of DREADDs-mediated inactivation. Left: Nissl stained sections and fluorescent images from individual animals used for the DREADDs experiments. In rat #169 and #170, recordings were performed from bilateral MEC and AAV (AAV8-hSyn-hM4Di-mCherry) was injected to the right RSC. Sagittal sections are shown for both Nissl-stained and fluorescent images. Positions of tetrode tracks are indicated by red circles. In rat #171 and #217, recordings were performed from the right RSC, and the AAV was injected to bilateral MEC. Coronal sections are shown for Nissl-stained images, and sagittal sections are shown for fluorescent images. Right two columns: the left plots show normalized firing rates of cells recorded from the virus injected site. The DREADDs agonist-21 was injected at the beginning of the recording sessions. Two red traces show representative cells that exhibited a significant reduction of firing rates after the injection (p < 0.05, Wilcoxon ranksum test for rate changes between 0-10 min and 30-40 min), and blue traces are the cells that were not significantly affected by the drug. The right plots show the probability density of the animal‟s running speed during random foraging in the open-field arena, before and after the drug injection. DREADDs-mediated inactivation did not significantly affect the animal‟s running speed (p > 0.05 in Friedman test). Each plot shows mean (solid lines) and s.e.m. (shaded).

## Methods

### Subjects

All experiments were approved by the local authorities (RP Darmstadt, protocol F126/1009) in concordance with the European Convention for the Protection of Vertebrate Animals used for Experimental and Other Scientific Purposes. Subjects were 10 male Long-Evans rats weighing 400 to 550 g (aged 3-5 months) at the start of the experiment. Rats were housed individually in Plexiglass cages (45 x 35 x 40cm; Tecniplast GR1800) and maintained a reversed 12-h light-dark cycle, with behavioural experiments performed during the dark phase. Animals were mildly food restricted with unlimited access to water, and kept at 85 to 90% of their free-feeding body-weight throughout the experiment. All rats had tetrodes located either unilaterally in RSC, of which six had a drive in the right hemisphere versus two animals in the left hemisphere, or bilaterally in MEC. Four rats were additionally injected bilaterally with an AAV encoding inhibitory DREADDs in either MEC or RSC. No statistical method was used to predetermine sample size, although the number of animals used here is similar to previous work.

### Surgery, virus injection and drive implantation

Anesthesia was induced by isoflurane (5% induction concentration, 0.5-2% maintenance adjusted according to physiological monitoring). For analgesia Buprenovet (Buprenorphine, 0.06 mg/mL; WdT) was administered by subcutaneous injection, followed by local intracutaneous application of either Bupivacain (Bupivacain hydrochloride, 0.5 mg/mL; Jenapharm) or Ropivacain (Ropivacain hydrochloride, 2 mg/mL; Fresenius Kabi) into the scalp. Rats were subsequently placed in a Kopf stereotaxic frame, and an incision was made in the scalp to expose the skull. After horizontal alignment several holes were drilled into the skull to place anchor screws, and craniotomies were made for microdrive implantation. The microdrive was fixed to the anchor screws with dental cement, while two screws above the cerebellum were connected to the electrode’s ground. All tetrodes were then positioned at 920 μm depth from the cortical surface. All animals received analgesics (Metacam, 2 mg/mL Meloxicam; Boehringer Ingelheim) and antibiotics (Baytril, 25 mg/mL Enrofloxacin; Bayer) for at least 5 days post-surgery.

For tetrode recordings, rats were unilaterally implanted with a hyperdrive that contained 28 individually adjustable tetrodes made from 17-μm polyimide-coated platinum-iridium (90-10%; California Fine Wire; plated with gold to impedances below 150 kΩ at 1 kHz). The tetrode bundle consisted of 30-gauge stainless steel cannulae, soldered together in a 14×2 rectangular shape for recordings of the entire RSC, 7×4 for anterior RSC, or two squared bundles for bilateral MEC. For RSC, tetrodes were implanted alongside the anteroposterior axis, starting at (AP) −2.5 mm posterior from bregma until −4 mm to −6.5 mm, (ML) 0.8 mm lateral from the midline, (DV) 1.0 mm below the dura, and at a 25° angle in a coronal plane pointing to the midline in order the get underneath the superior sagittal sinus. For MEC, tetrodes were implanted at 4.5 mm lateral of the midline, 0.2 mm anterior to the transverse sinus, at an angle of 15 degrees in a sagittal plane with the tips pointing to the anterior direction. Experiments began at least 1 week post-surgery to allow the animals to recover.

For DREADDs experiments, an AAV8-hSyn-hM4Di-mCherry (a gift from Bryan Roth; Addgene viral prep #44362-AAV8) was injected with an infusion rate of 100 nL/min using a 10 μl NanoFil syringe and a 33-gauge bevelled metal needle (World Precision Instruments). After injection was completed the needle was left in place for 10 min. The virus was injected at two sites for each bilateral MEC (500 nL each at the depth of 2.5 mm and 3.5 mm from the cortical surface, 4.5 mm lateral to the midline, 0.2 mm anterior to the transverse sinus at an angle of 20° in a sagittal plane with the needle pointing to the anterior direction), or 4 sites along the anteroposterior axis for each bilateral RSC (500 nL each at AP 2.5, 3.5, 4.5, 5.5 mm, 0.8 mm lateral to the midline, at an angle of 25° in a coronal plane pointing to the midline). Flow was controlled with a Micro4 microsyringe pump controller. A small microdrive (Axona) connected to 4 wire tetrodes was additionally implanted nearby the injection site to evaluate the effects of the manipulation. Virus injection was performed in the same surgery as electrode implantation, and recordings began at least three weeks post-surgery to allow time for the virus to express.

### Spike sorting and cell classification

All main analyses and data processing steps were performed in MatLab (MathWorks). Neural signals were acquired and amplified using two 64-channel RHD2164 headstages (Intan technologies), combined with an OpenEphys acquisition system, sampling data at 15 kHz. Neuronal spikes were detected by passing a digitally band-pass filtered LFP (0.6-6 kHz) through the ‘Kilosort’ algorithm to isolate individual spikes and assign them to separate clusters based on waveform properties (https://github.com/cortex-lab/KiloSort) (Pachitariu, Steinmetz, Kadir, Carandini, & Harris, 2016). Clusters were manually checked and adjusted in autocorrelograms and for waveform characteristics in principal component space to obtain well-isolated single units, discarding any multi-unit or noise clusters.

### RSC border cells

We applied a novel template-matching procedure to classify RSC neurons as border cells using the Earth Mover’s Distance (EMD), a distance metric from the mathematical theory of optimal transport (Hitchcock, 1941; Rubner et al., 1998). First, the animal’s spatial position occupancy was divided into 4×4 cm spatial bins, and the firing rate in each position bin was calculated by dividing the number of spikes with the amount of time spent there. The resulting ratemap was smoothed by applying a 2D Gaussian filter (width of 1 bin), and converted to a probability distribution by taking unit weight. We then calculated the Earth Mover’s Distance relative to a “border template” using a MatLab implementation of the fastEMD algorithm (https://github.com/dkoslicki/EMDeBruijn) (Pele & Werman, 2008, 2009). This border template consisted of a 25×25 matrix with each bin’s value set to 0, except the outer ring bins with a value of 1, smoothed with the same Gaussian kernel and converted to unit weight. Several additional templates were constructed to assess the effects of behavioural and neural manipulations (see Fig. 2, 3), adding additional weight in the location of placed objects/walls, or removing it in the absence of an outer wall. The EMD distance between a ratemap and a template represents the minimal cost that must be paid to transform one distribution into another, and is thus a normalized metric of dissimilarity (Grossberger et al., 2018).

To assess whether a cell’s ratemap was significantly similar to the border template, we computed a null distribution to compare against using Monte Carlo simulations. We performed 32.000 permutations of a shuffling procedure, and for each iteration we randomly sampled a spike-train from the data, time-shifted this vector along the animal’s recorded trajectory by a random interval of at least 4 seconds and less than the total trial length, wrapping any excess at the and back to the beginning. We then used this shifted data to compute a ratemap and calculated the EMD distance relative to the border template. Criteria for border cell classification was an EMD dissimilarity score below the 1^st^ percentile of this null distribution in all regular sessions, and an average firing rate of at least 0.5 Hz (see Fig. 1d, 1e).

### MEC border cells

To compare classification results with a related metric we computed the original border score for each cell (Solstad et al., 2008). We first estimated a cell’s firing field by isolating a continuous region of at least 200 cm^2^ and a maximum of 70% of the arena surface where the firing rate was above 30% of the peak firing rate. This was an iterative search until all fields with the above criteria were identified. We next computed the border score, b, for each wall separately:

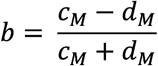

where c_m_ was defined as the maximum coverage of any single field over the wall and d_m_ the mean firing distance, calculated as the average distance to the nearest wall over all bins covered by the field. This was done separately for each of the four walls out of which the maximum score was selected. Cells recorded in MEC were classified as border cells whenever their border score was above the threshold of 0.5 (corresponding to the 99.3 percentile of scores generated from randomly time-shifted spikes) for either of the two recorded sessions, and had an average firing rate of at least 0.5 Hz.

### Head direction cells

The rat’s head direction was calculated based on the relative x/y-position of two light emitting diodes (LEDs), corrected for an off-set in placement of the LED’s relative to the animal’s true head direction. For each cell the mean vector length (MVL) and direction (MVD) was calculated by computing the circular mean and direction from a vector that contained the head direction of the animal at spike timings in unit space. A cell was classified as a head direction cell when its MVL was greater than the 95^th^ percentile of a null distribution obtained by 1000-fold Monte Carlo simulations with randomly time-shifted spike trains.

### Border rate maps

Locations of walls were estimated based on the most extreme values of the position of the animal. The animal’s distance to the wall was computed for each of the four walls separately by taking the difference between the wall’s location and the animal’s position in the respective x or y-dimension, and selecting the lowest value at each time point. The direction of this wall relative to the animal’s direction was computed by calculating the angle difference between the animal’s true heading direction and a vector pointing directly towards the wall (e.g. relative to an angle of 0° for the east wall, 90° for the north wall etc.). Because 0° corresponds with the ‘east’ side in angular polar plots, this data was further shifted by 90° to align the front of the animal with the ‘north’ part in border maps (see Fig. 4c) to improve visual interpretation of the results.

Firing rate in these body-centric border coordinates was calculated by dividing the animal’s occupancy in these coordinates into 4 cm distance bins and 20° angle bins. The number of spikes in each bin was then divided by the time spent there, further smoothed using a 2-D Gaussian kernel (1 bin width), similar to how spatial rate maps are computed. A cell’s preferred direction and distance was obtained by finding the bin with maximal firing rate, and selecting the bin’s corresponding distance and angle values. For visualization purposes only this matrix was transformed into a circular diagram shown in Fig 4.

### Decoding analysis

For decoding of wall distance from the activity of border cells in RSC and MEC, the optimal wall with maximum coverage by firing fields was chosen for individual cells (the same procedure as used in border-score calculations (Solstad et al., 2008)). To determine the optimal head direction to the selected wall for individual border cells, we searched for a range of head directions (60-degree range in 5-degree steps) that gave the maximum mean firing rate of the cell when the animal was within 20 cm of the wall. We then focused on neural activity when the animal was at this optimal head direction and in the range of wall distances from 0 to 50 cm at 10 cm steps (5 ranges in total), but excluding timepoints where the animal was within 25 cm of other walls to avoid their potential influence. All of the incidents when the animal was in each of the 5 wall-distance ranges were equally divided into 20 segments in time, and mean firing rates of individual border cells in the 20 segments were assembled together across recording sessions. To implement a decoding analysis, 20 cells were randomly chosen, and the order of 20 segments was randomly shuffled for each cell, such that the data in each segment is a collection of firing rates from 20 border cells across various time points of behaviours when the animal was in a particular distance range to the wall. Ensemble firing rates of border cells in one of the segments were selected as a test dataset, and the rest of the data were used to train a support vector machine (using a MATLAB package LibSVM with a linear function (Chang & Lin, 2011)). Trained weights were then applied to the activity of border cells in the test dataset to estimate the animal‟s distance to the wall, which was repeated for all segments to be tested (leave-one-out cross-validation), giving a representative decoding performance for the selected population of cells. This procedure was repeated for different cell pairs for 1000 times to estimate a statistical distribution of decoding performance (bootstrap resampling method).

### Behavioural methods

Data was collected over a total of 30-120 min per day while rats foraged for food (chocolate cereal) in a squared open field arena, either 100×100 cm or 120×120 cm in size. Each session consisted of 10-15 min of free exploration in the arena, separated by 5-10 min of resting time on a pedestal. No curtains surrounded the recording arena, with the exception of the rotation and darkness experiments where all distal cues were blocked completely. The surface of the arena was elevated 50 cm above the ground, and was enclosed by three black and one white wall with a 50 cm height that were positioned with consistent orientation in the room for all animals. The experimental set-up was extensively cleaned with a 70% ethanol solution in between every recording session to eliminate any odours.

Behavioural manipulation experiments always followed the same protocol of A-B-B-A’, where A is a regular session, and the manipulation was performed in B. This allowed for a recovery phase after the manipulation in the final session A’. The only exception was the drop-edge experiment (Fig. 3e) where the animal had limited motivation; so to ensure good coverage of the arena we reduced the protocol to A-B-A’. All changes to the maze were made in between the first and second session while the animal was resting on a pedestal. For the added wall manipulation (Fig. 2a), an additional black wall (50 cm length x 50 cm height x 1 cm width) was placed in the maze, protruding from one outer wall at half-length towards the centre. For the added object manipulation (Fig. 2f) a circular, non-climbable aluminium object (10 cm diameter x 50 cm height) was placed off-centre 40 cm away from the north and west walls. For the DREADDs-mediated manipulation experiments, animals were injected with agonist-21 (DREADDs agonist 21 dihydrochloride, 3.52 mg/mL [10 mM]; Hellobio) subcutaneously after the first recording session, followed by at least 30 min waiting time to allow the drug to reach the brain and take effect before starting the next recording session.

The animal’s position and head direction were obtained by tracking two LEDs on the headstage at 25 Hz and recording under dimly lit conditions. For darkness sessions, we switched to an infra-red OptiTrack camera system (Natural Points Inc.). Six Flex 3 cameras were place around the experimental set-up that recorded the location of three reflective markers in an asymmetric frame attached to the headstage. Position and direction data were acquired and processed using Motive 2.0 software.

### Histological procedures

Once the experiment was completed, animals were deeply anesthetized by sodium pentobarbital and perfused intracardially with saline, followed by 10% formalin solution. Brains were extracted and fixed in formalin for at least 72 hours at 6° C temperature. Frozen coronal sections were cut (50 μm) and stained using cresyl violet and mounted on glass slides. Electrode tips were identified by comparison across adjacent sections, with the location of recorded cells estimated by backward measurement from the most ventral tip of the tetrode tracks.

### Statistical procedures

All statistical tests were two-sided and non-parametric, unless stated otherwise. Error bars in all figures represent standard error of the mean (S.E.M.). All values mentioned in text are medians ± S.E.M.

## Acknowledgements

We thank Martin Vinck for suggesting the approach with the Earth Mover Distance (EMD) and providing initial software for analysis. We also thank Diogo Santos-Pata for discussion and comments related to the manuscript. This work was supported by the Max Planck Society, the Behrens-Weise Foundation, and the European Research Council („NavigationCircuits‟ Grant Agreement no. 714642)

## Author Contributions

J.B.G.v.W. and H.T.I. designed the experiment. J.B.G.v.W. performed all experiments and analyses, except for the cue-rotation experiment and DREADDs experiment, which were performed by S.S.B. and H.T.I. respectively. J.B.G.v.W. and H.T.I. wrote the manuscript.

